# A sperm-enriched 5’fragment of tRNA-Valine regulates preimplantation embryonic transcriptome and development

**DOI:** 10.1101/2024.08.08.607197

**Authors:** Simeiyun Liu, Andrew D. Holmes, Sol Katzman, Upasna Sharma

## Abstract

Sperm small RNAs have been implicated in intergenerational epigenetic inheritance of paternal environmental effects; however, their biogenesis and functions remain poorly understood. We previously identified a 5’ fragment of tRNA-Valine-CAC-2 (tRFValCAC) as one of the most abundant small RNA in mature sperm. tRFValCAC is specifically enriched in sperm during post-testicular maturation in the epididymis, and we found that it is delivered to sperm from epididymis epithelial cells via extracellular vesicles. Here, we investigated the mechanistic basis of tRFValCAC delivery to sperm and its functions in the early embryo. We show that tRFValCAC interacts with an RNA binding protein, heterogeneous nuclear ribonucleoprotein A/B (hnRNPAB), in the epididymis, and this interaction regulates the sorting and packing of tRFValCAC into extracellular vesicles. In the embryo, we found that tRFValCAC regulates early embryonic mRNA processing and splicing. Inhibition of tRFValCAC in preimplantation embryos altered the transcript abundance of genes involved in RNA splicing and mRNA processing. Importantly, tRFValCAC-inhibited embryos showed altered mRNA splicing, including alternative splicing of various splicing factors and genes important for proper preimplantation embryonic development. Finally, we find that inhibition of tRFValCAC in zygotes delayed preimplantation embryonic development. Together, our results reveal a novel function of a sperm-enriched tRF in regulating alternating splicing and preimplantation embryonic development and shed light on the mechanism of sperm small RNA-mediated epigenetic inheritance.

## INTRODUCTION

Small RNAs play key roles in multiple, well-established epigenetic inheritance paradigms, such as RNA interference in *C. elegans* and paramutation in maize [1, 2]. A wide variety of distinct species of small RNAs have been described, including well-studied small RNAs, such as microRNAs, endo-siRNAs, and gamete-enriched piRNAs, as well as relatively understudied small RNAs, such as cleavage products of tRNAs, known as tRNA fragments (tRFs) or tRNA-derived small RNAs (tsRNAs). Recently, tRNA-derived RNA (tDR) has been proposed as a uniform naming system for tRNA fragments that builds on the existing naming system of full-length tRNAs [3]. For this study, the cleavage products of tRNAs will be described as tRFs to be consistent with previous studies relevant to this work [4, 5]. tRFs are a recently discovered class of small RNAs widespread in most organisms [6–9] and can be derived from either 5’, 3’, or the middle of the full-length tRNA molecules. Across different organisms and cell types, tRFs have been proposed to play diverse biological roles, including global and transcript-specific translation inhibition [10–13], ribosome biogenesis [14–17], repression of mRNAs via either 3’UTR targeting or Dicer-dependent or -independent silencing [18–24], alteration of chromatin accessibility [4, 25, 26], regulation of apoptosis [27], modulation of cell proliferation [18], and regulation of retrotransposons [26, 28].

tRFs are one of the most abundant classes of small RNAs in mature mammalian sperm [4, 29]. Intriguingly, the small RNA composition of sperm undergoes significant alterations during post-testicular maturation within the epididymis, an elongated tubular organ that links the testis to the vas deferens in the male reproductive system and is required for sperm motility and fertility. We previously reported that while testicular spermatozoa are highly enriched in gamete-specific piRNAs, the mature sperm in the epididymis have a high abundance of tRFs [4, 5]. tRFs are also highly abundant in the somatic epididymis epithelial cells, and a subset of these tRFs can be delivered to sperm via extracellular vesicles (EVs) secreted from epididymis epithelial cells, known as epididymosomes [4, 5]. The mechanism of this soma-to-germline delivery of small RNAs via extracellular vesicles remains unknown.

Importantly, sperm tRFs are modulated by environmental conditions and can influence offspring phenotypes. For instance, exposure of male mice to dietary alterations (such as a high-fat, low-protein, western or high-sugar diet), inflammation, gut-microbiota depletion, aging, ethanol, and other toxicants (DDT, phthalates, methotrexate, and opioids) resulted in altered levels of tRFs in sperm of mice [4, 30–37], fishes [38] and men [39, 40]. Moreover, injection of bulk tRFs purified from the sperm of exposed males or environment-sensitive synthetic tRFs into control zygotes generated adult offspring with altered phenotypes [30, 33, 34, 41]. However, the specific roles sperm-enriched tRFs play in regulating offspring phenotypes remains to be elucidated. Understanding the biogenesis and functions of sperm tRFs is crucial to elucidate their role in intergenerational epigenetic inheritance. Moreover, as relatively immature sperm from testicular biopsies are used to fertilize oocytes in some cases of assisted reproduction in humans, it is vital to understand the functional consequences of mature sperm small RNA payload.

A 5’ fragment of tRNA-Valine-CAC-2 (tRFValCAC) is among the most abundant small RNAs that change dramatically in abundance as sperm mature in the epididymis [4]. Our previous studies revealed that tRFValCAC is enriched in epididymosomes and is one of the small RNAs delivered to sperm via epididymosomes [4]. Here, we set out to investigate the biogenesis and functions of sperm-enriched tRFValCAC. We found that an RNA-binding protein, hnRNPAB, directly interacts with tRFValCAC in the epididymis. Knockdown of hnRNPAB in epididymis epithelial cells led to reduced levels of tRFValCAC in the EVs secreted from those cells, implicating hnRNPAB in sorting of tRFValCAC into epididymosomes and, consequently, sperm delivery. Investigation of tRFValCAC function in the early embryo revealed that inhibition of tRFValCAC using an antisense locked Nucleic Acid (LNA) containing oligo (Val_LNA) altered transcript abundance of numerous genes at the 2-cell stage of preimplantation embryos, including those involved in RNA splicing and mRNA processing. Global alternative splicing analysis of Val_LNA embryos revealed altered splicing compared to control embryos. Finally, Val_LNA embryos had delayed preimplantation embryonic development, with a significantly lower percentage of Val_LNA embryos reaching the blastocyst stage compared to control embryos. These studies reveal a novel function of sperm-enriched tRFValCAC in regulating early embryonic gene expression and development.

## RESULTS

### tRFValCAC interacts with hnRNPAB in the epididymis, and this interaction regulates its sorting to EVs

During epididymal maturation, sperm first enter the proximal caput epididymis; from there, they progress to the corpus, and finally, mature sperm are stored in the cauda epididymis (**Figure 1A**). We previously found that tRFValCAC is one of the most abundant small RNA in mature sperm and that tRFValCAC is delivered to sperm from the epididymis epithelial cells via epididymosomes [4]. Here, we aimed to elucidate the mechanism of tRFValCAC enrichment in sperm during epididymal maturation. RNA-binding proteins oversee every facet of RNA existence, encompassing its creation, alteration, movement within cells, biological roles, and eventual degradation [42]. To identify tRFValCAC interacting proteins in the epididymis, we used a biotinylated tRFValCAC mimic to pull down proteins that interact with tRFValCAC from the epididymal cellular extract (**Table S1**). 16 proteins were significantly enriched in tRFValCAC pull-downs relative to the scrambled control (**Table S2**), and their biological functions and protein-protein interaction networks were analyzed using the PANTHER classification system. Gene Ontology analysis showed that 11 out of 16 proteins were enriched in the RNA binding category, and 5 of those proteins were overrepresented in the mRNA metabolic processes (**Table S2**). From the top candidate interacting proteins, we focused on hnRNPAB (**Figure 1B**), which belongs to the heterogeneous nuclear ribonucleoprotein (hnRNPs) family of proteins that have been implicated in gene regulation and miRNA sorting and packaging into EVs [43–46]. Moreover, hnRNPF and hnRNPH were recently identified as direct binding partners of a 5’ fragment of tRNA-Glycine-GCC [26].

**Figure 1:**
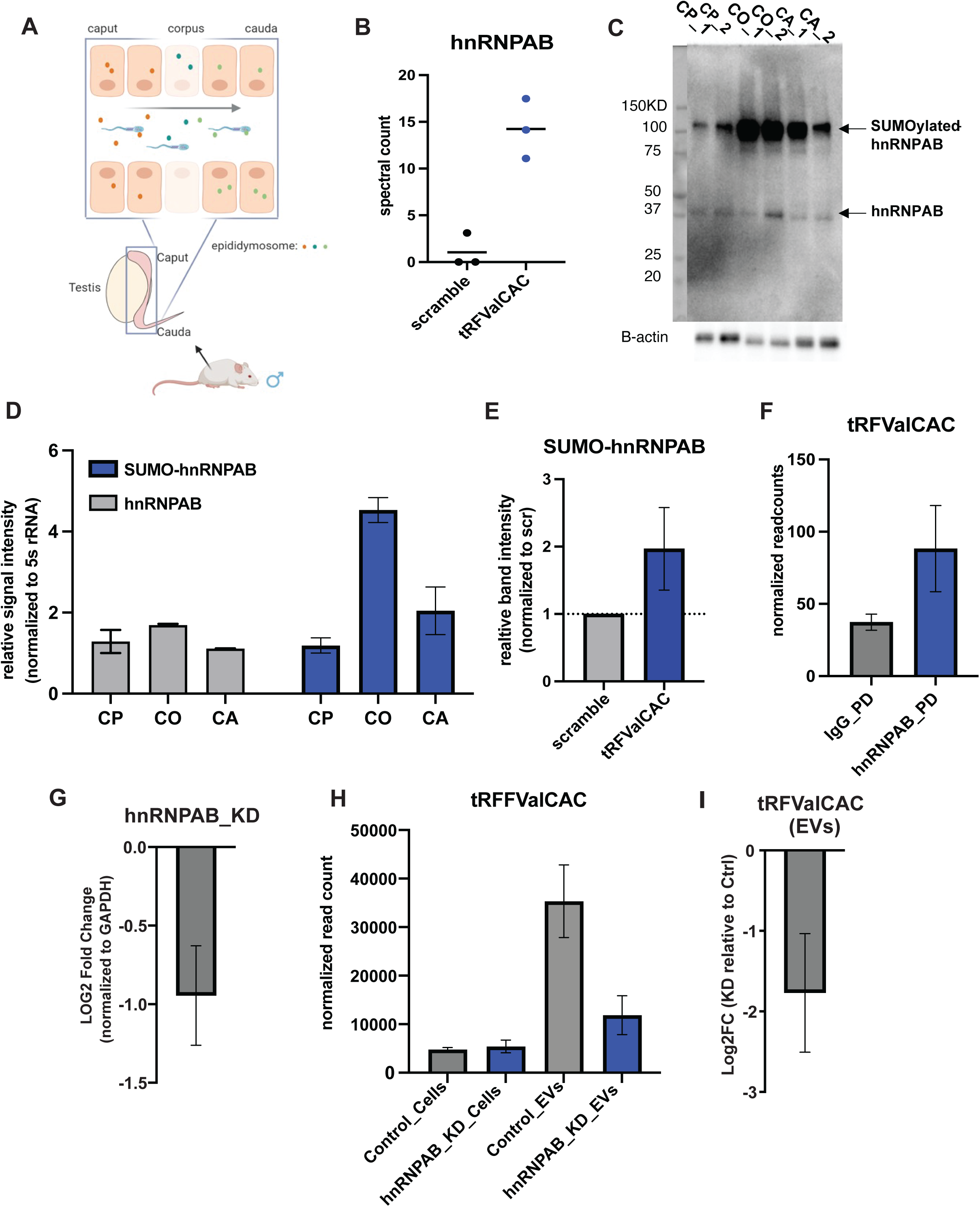
hnRNPAB interacts with tRFValCAC in the epididymis and regulates its sorting into extracellular vesicles. **A)** Schematic depicting the direction of sperm movement through the caput, corpus, and cauda epididymis. Epididymosomes secreted from the epididymis epithelial cells fuse with sperm to deliver proteins and small RNAs. **B**) Spectral counts of hnRNPAB protein in LC-MS data of proteins pulled down using a biotinylated tRFValCAC synthetic oligo. hnRNPAB is significantly enriched in tRFValCAC pull down compared to scramble control (*p* value=0.01). **C)** Western blot analysis of total protein isolated from caput (CP), corpus (CO), and cauda (CA) epididymides using anti-hnRNPAB. The top band is higher than the expected molecular weight of hnRNPAB (∼37KD). It represents a SUMOylated form of hnRNPAB (based on Western blot analysis using an anti-SUMO antibody; see Supplementary Figure S1A). The lower band is the unmodified hnRNPAB and is of expected size. Beta-actin is used as a loading control. The leftmost gel image is the image of the Kaleidoscope Precision Plus Protein Standard ladder from the same gel imaged under a trans-white light setting for the estimation of the molecular weight of the proteins. **D)** Quantification of SUMO-hnRNPAB and hnRNPAB bands from (C) relative to loading to control beta-actin (n=2). **E)** Western blot analysis using the anti-hnRNPAB antibody of proteins pulled down with tRFValCAC oligo (n=6 replicates). The graph shows the quantification of hnRNPAB band intensity in tRFValCAC pulldown relative to Scramble control oligo pulldown. The Western blot image is included in supplementary Figure S1C. As SUMO-hnRNPAB is the main protein detected in the Western blot, that band was used for quantification in tRFValCAC pull-down relative to the band of the same size in scramble control. **F)** Normalized read counts of tRFValCAC (5’ fragment of tRNA-ValCAC-2) in small RNA sequencing libraries generated from RNA immunoprecipitated using beads conjugated with anti-hnRNPAB antibody (hnRNPAB-PD) or IgG bead control (IgG-PD) (n=2 replicates). **G)** Knockdown of hnRNPAB in DC2 cells using siRNAs pool targeting *hnrnpab* transcript. The bar graph shows a log2 fold change in hnRNPAB transcript abundance relative to B-actin control. **H)** Normalized read counts of tRFValCAC in small RNA sequencing data generated from hnRNPAB knockdown or control DC2 cells and their EVs. DC2_C_CELL and DC2_KD_CELL represent RNA of control (C) and hnRNPAB knockdown (KD) DC2 cells. DC2_C_EV and DC2_KD_EV represent RNA of EVs purified from media of control and hnRNPAB knockdown cells, respectively. **I)** Log2 fold change of tRFValCAC in knockdown (KD) relative to control (C) EVs. Graphs represent data as mean ± SEM.

We examined the expression pattern of hnRNPAB across different segments of the epididymis (caput, corpus, and cauda epididymis). We found that hnRNPAB is expressed in the epididymis in both SUMOylated (small ubiquitin-like modifier) and unmodified forms, with SUMOylated (SUMO-hnRNPAB) form having a higher molecular weight as observed in Western blot analysis (**Figure 1C and S1A**). Notably, although the levels of unmodified hnRNPAB remained uniform throughout different segments of the epididymis, there were segment-specific differences in the levels of the SUMO-hnRNPAB. Cauda epididymis had a higher abundance of SUMO-hnRNPAB than caput epididymis, with peak levels in the corpus epididymis (**Figure 1C-D**). Moreover, we also detected SUMO-hnRNPAB in epididymosomes (**Figure S1B**). Having detected hnRNPAB in the epididymis, we further examined the interaction between tRFValCAC and hnRNPAB. Western blot analysis of tRFValCAC pull-down proteins also revealed enrichment of hnRNPAB in tRFValCAC pull-downs compared to the scramble control (**Figure 1E, Figure S1C**).

Consistent with the high abundance of SUMO-hnRNPAB in the epididymis, the SUMOylated form of hnRNPAB was detected in the pull-downs (**Figure S1C**). In addition, immunoprecipitation of hnRNPAB-interacting RNAs showed enrichment of tRFValCAC in the hnRNPAB pull-downs compared to IgG control pull-downs (**Figure 1F, Figure S1D**). Together, these data demonstrated that hnRNPAB binds to tRFValCAC in the epididymis.

Due to low amounts of cellular extract, we could not directly assay tRFValCAC-hnRNPAB interaction in epididymosomes. Nonetheless, as hnRNPAB is detectable in epididymosomes (**Figure S1B**) and hnRNPA2B1 has been implicated in miRNA sorting into EVs [43], we hypothesized that hnRNPAB plays a role in tRFValCAC sorting into epididymosomes. We utilized an epididymis epithelial cell line derived from the distal caput epididymis (DC2) [47] to examine the role of hnRNPAB in tRFValCAC sorting into epididymosomes. First, we confirmed tRFValCAC-hnRNPAB interaction in DC2 cells. Western blot analysis of proteins pulled-downs with tRFValCAC oligo from DC2 cellular extract showed significant enrichment of hnRNPAB in the tRFValCAC pull-downs compared to the scramble control (**Figure S1E-F**). However, SUMO-hnRNPAB was not the dominant form of this protein in DC2 cells, demonstrating differential regulation of hnRNPAB in cultured cells. Next, we evaluated the enrichment of tRFValCAC in EVs secreted by DC2 cells. Mimicking selective sorting of tRFValCAC into epididymosomes *in vivo* [4, 5], sequencing profiles of small RNA content in DC2 cells and their EVs revealed a significant enrichment of tRFValCAC in EVs relative to epithelial cells (**Figure S1G**). These results established DC2 cells as a suitable *in vitro* system for studying tRFValCAC sorting and packing into epididymosomes.

To examine the role of hnRNPAB-tRFValCAC interaction in regulating tRFValCAC sorting into EVs, we knocked down hnRNPAB in DC2 cells using a pool of siRNAs (**Figure 1G**). Next, we examined the abundance of tRFValCAC in the knockdown cells and EVs secreted from those cells. Interestingly, the knocking down of hnRNPAB resulted in reduced levels of tRFValCAC in EVs without significant changes in cellular levels (**Figure 1H-I**). These results demonstrate that hnRNPAB regulates tRFValCAC levels in the EVs secreted from epididymis epithelial cells. This process was not dependent on SUMOylation of the protein, as only an unmodified form of hnRNPAB was detected in DC2 cells. SUMOylation of hnRNPAB potentially regulates other functions of the tRFValCAC-hnRNPAB complex *in vivo*. Together, these data identified RNA binding protein hnRNPAB as a binding partner for tRFValCAC in the epididymis, with a potential role in regulating the sorting of tRFValCAC into epididymosomes and, thereby, its delivery to sperm [5].

### tRFValCAC regulates the transcriptome of preimplantation embryos

Sperm small RNAs are transmitted to the embryo at fertilization and, thus, function in paternal epigenetic inheritance [48–51]. Therefore, we examined the regulatory roles of tRFValCAC in early preimplantation embryos. We inhibited tRFValCAC using an antisense locked-nucleic acid (LNA) containing oligo targeting tRFValCAC (Val_LNA) [52]. Val_LNA successfully inhibited tRFValCAC (**Figure S2A**) and did not affect the levels of full-length mature tRNA-Val-CAC-2 (**Figure S2B**) in mouse embryonic stem cells, demonstrating its specificity to tRFValCAC. Embryos were generated using *in-vitro* fertilization (IVF) and injected with either Val_LNA or a control LNA oligo (Ctrl_LNA) (**Figure 2A**). Embryos were harvested at the late 2-cell stage post zygotic genome activation and processed for single-embryo mRNA sequencing. Principal component analysis (PCA) showed separate clustering of individual embryos from the two experimental conditions (**Figure 2B**). Differential gene expression analysis revealed 505 differentially expressed genes between Val_LNA and Ctrl_LNA embryos (*padj* value<0.05, Log2 Fold change ≥1); of these, 311 genes were downregulated and 194 upregulated (**Figure 2C and Table S3**). Gene ontology (GO) enrichment analysis of overrepresented pathways in differentially expressed genes showed an enrichment of genes involved in chromosome organization, cell cycle, and mRNA processing and splicing (**Figure 2D**). For instance, genes encoding for histone modifying enzymes were dysregulated, including histone demethylase *Kdm2b* and *Kdm4a*, which were down- and upregulated, respectively (**Figure 2E**). Moreover, cell cycle regulator cyclin E1 (*Ccne1*) was downregulated, and the histone H2B gene (*H2bc4*) was upregulated (**Figure 2E**). Consistent with these observations, histone genes were overall upregulated in Val_LNA embryos compared to all other genes (**Figure 2F**).

**Figure 2:**
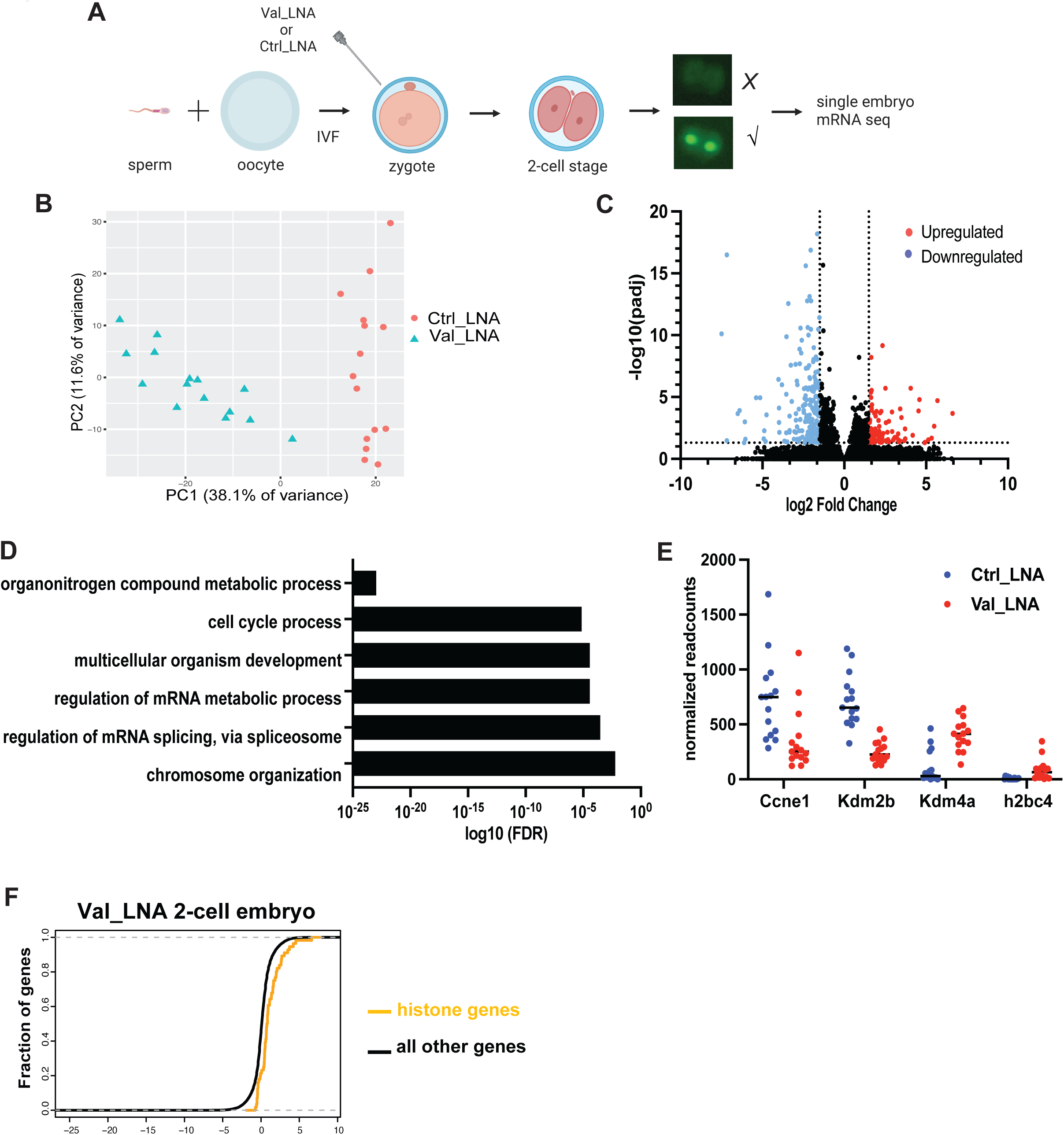
tRFValCAC regulates early embryonic transcriptome. **A)** Embryos were generated using in-vitro fertilization (IVF) and injected with histone H3.3 fused GFP mRNA plus either LNA oligo targeting tRFValCAC (Val_LNA) or a control LNA oligo (Ctrl_LNA). H3.3-GFP was added to allow the selection of successfully injected embryos using GFP fluorescence. To control for total RNA molecules per embryo, we injected the same amount of total RNA in both conditions. Further, the Ctrl_LNA oligo has the same number of LNA nucleotides as Val_LNA, thus controlling for any cellular toxicity of the LNAs. As shown in Figure S2A-B, Val_LNA successfully inhibited tRFValCAC and did not affect full-length mature tRNA-Val-CAC-2 levels in mouse embryonic stem cells. **B)** PCA analysis of transcriptomic data from Ctrl_LNA and Val_LNA injected embryos (n=15 for each condition). **C)** Volcano plot representing differentially expressed genes in Val_LNA embryo relative to Ctrl_LNA embryos (genes that showed Log2 fold change of 1.5 or higher with *padj* value <0.05). See Table S3 for the full list of differentially expressed genes. **D)** Gene ontology (GO) analysis of biological processes overrepresented in significantly differentially expressed genes. **E)** Normalized read counts of transcripts of genes involved in chromatin regulation and cell cycle. Each data point represents a single embryo. *padj* values from DESeq2 analysis: *Ccne1*- 0.02, *Kdm2b*- 4.1×10^-12^, *Kdm4a*- 0.03, *h2bc4*- 0.0004. **F)** Cumulative distribution plots showing transcript abundance of histone genes and all other genes in Val_LNA relative to Ctrl_LNA embryos.

tRFs carry modified residues important for their stability and function [13, 53, 54]. As the modification status of tRFValCAC is currently unknown, we could not synthesize a tRFValCAC oligo that mimics the endogenous tRFValCAC to study its gain-of-function effects in embryos. Therefore, we aimed to induce the generation of endogenous tRFValCAC using mouse embryonic stem cells (mESCs) that serve as a mechanistically tractable *in vitro* system with close physiological resemblance to embryonic conditions. A previous study revealed that retinoic acid (RA) treatment induces the generation of 5’ fragments of tRNA-Valine-CAC in mouse embryonic stem cells [55]. Leveraging this knowledge, we used RA treatment to induce tRFValCAC in mESCs (**Figure S2C)** and examined changes in the abundance of tRFValCAC-target transcripts identified in mouse embryos. An increase of tRFValCAC is predicted to induce changes in gene expression that are opposite to those observed with tRFValCAC inhibition in embryos. Indeed, RA-induced increase of tRFValCAC in mESCs resulted in downregulation of *H2bc4* and *Kdm4a* (**Figure S2D**). Together, these data demonstrate that inhibition of tRFValCAC results in altered transcript abundance of genes involved in cell-cycle and chromatin organization. Given the critical roles of these processes for proper embryonic development, these changes in embryonic transcriptome potentially influence preimplantation embryonic development (see below).

### tRFValCAC regulates RNA processing and splicing in preimplantation embryos

In addition to cell cycle and chromosome organization, GO analysis revealed that genes involved in RNA processing, splicing, and spliceosome were overrepresented in the differentially expressed gene in Val_LNA embryos (**Figure 2E and S3A**). Indeed, numerous genes involved in RNA processing and splicing were misregulated in Val_LNA embryos, including the downregulation of splicing factors *Srsf10* and *Srrm2* (**Figure 3A**). Consistent with changes in Val_LNA embryos, induction of tRFValCAC using RA treatment resulted in an upregulation of *Srsf10* and *Srrm2* transcripts in mESCs (**Figure S3B**). Next, we performed global alternative splicing analysis to determine if alterations in transcript abundance of RNA splicing regulators in Val_LNA embryos led to splicing defects in these embryos. As single embryo mRNA-sequencing data is incompatible with alternative splicing analysis due to low read counts per embryo, we pooled sequencing reads from 4-5 embryos to generate pseudo-replicates and performed RNA splicing analysis using junctionCounts (**Methods**).

**Figure 3:**
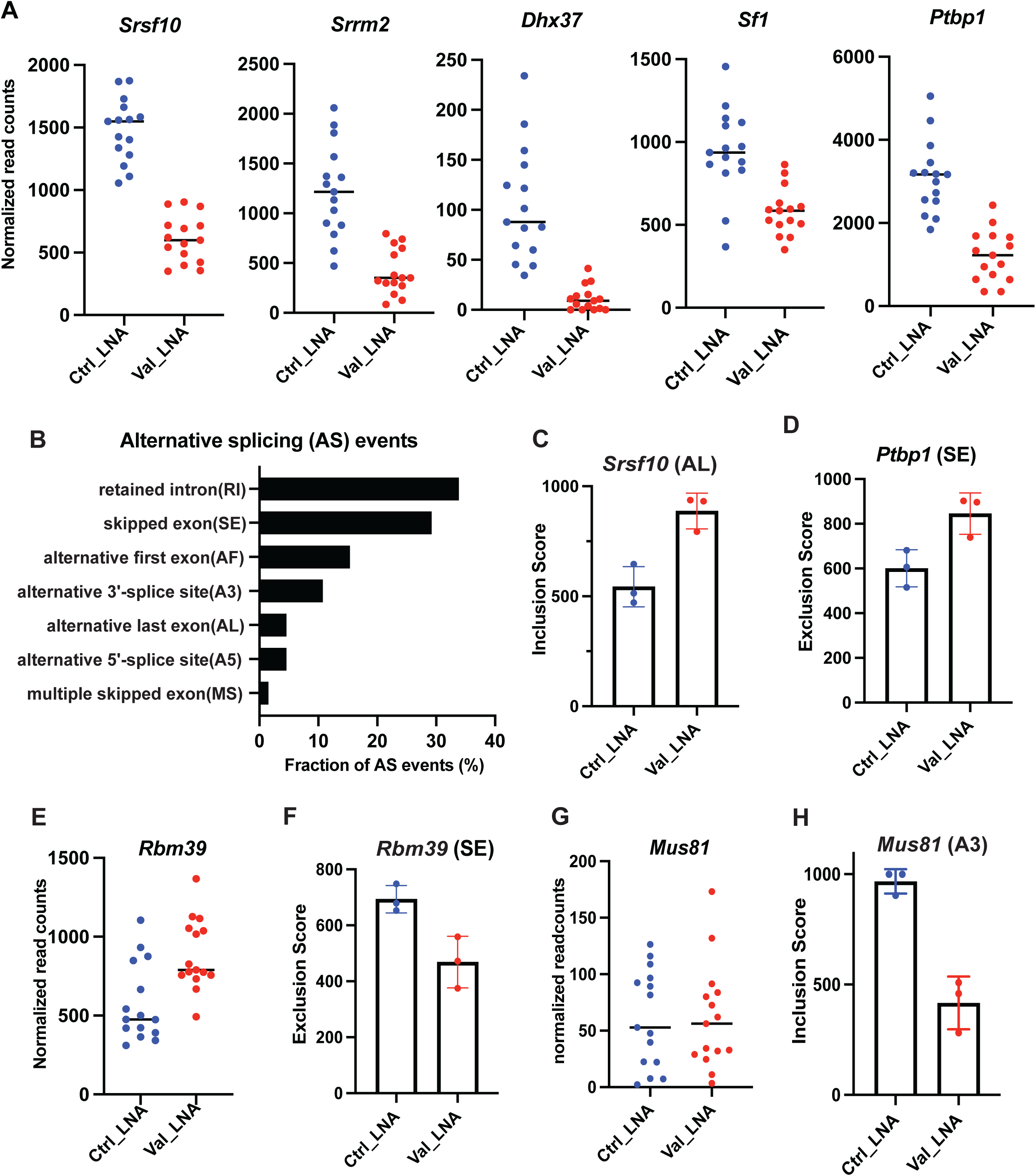
Defective alternative splicing in tRFValCAC-inhibited embryos. **A)** Normalized read counts of a subset of RNA processing and splicing factors that are significantly differentially expressed in Val_LNA embryos relative to Ctrl_LNA embryos (DESeq2 *padj* values: *Srsf10* 1.76×10^-16^, *Srrm2* 1.06×10^-6^, *Dhx37* 8.00 × 10^-8^, *Sf1* 6.02 × 10^-4^, and *Ptbp1* 1.76 × 10^-6^). Each data point represents a single embryo. **B)** The plot shows the number of significant alternative splicing events for each event type (△PSI > 0.15 and pairSum 7 or higher). The number of significant events for each event type is calculated as a fraction of all significant events in Val_LNA embryos relative to Ctrl_LNA embryos (total significant events = 65). Sequencing reads from 5-4 embryos were pooled to generate pseudo-replicates (n=3). **C)** Inclusion score of alternate last exon 6 (AL) in *Srsf10* transcripts in Val_LNA and Ctrl_LNA embryos. A browser track visualization of this alternative splicing event is shown in Supplementary Figure S3D. **D)** Exclusion score of the exon 8 in *Ptbp1* transcripts. A browser track visualization of this alternative splicing event is shown in Supplementary Figure S3C. **E)** Normalized read counts of *Rbm39*. Val_LNA embryos displayed a higher abundance of *Rbm39* transcript relative to Crtl_LNA embryos (DESeq2 *padj* value 0.02). **F)** Exclusion score of an exon skipping event in *Rbm39* transcript in Val_LNA and Ctrl_LNA embryos. A browser track visualization of this alternative splicing event is shown in Supplementary Figure S3E. **G)** *Mus81* transcript abundance in Val_LNA and Ctrl_LNA embryos (DESeq2 *padj* value 0.98). **H)** Inclusion score of an alternative 3’ splice site in *Mus81* transcript. A browser track visualization of this alternative splicing event is shown in Supplementary Figure S3F.

A total of 65 significant alternative splicing events were identified (ΔPSI >0.15, pairSum ≥ 7), with approximately one-third being retained introns and another one-third being skipped exons (**Figure 3B and Table S4**). Interestingly, we detected significant alternative splicing events in splicing regulators *Srsf10*, *Ptbp1*, and *Rbm39*. In Val_LNA embryos, an alternate last exon (AL) was included at a higher level in *Srsf10* transcripts relative to Ctrl_LNA embryos **(Figure 3C and S3D).** *Ptbp1* transcripts displayed exon skipping events with higher levels of exclusion of exon 8 in Val_LNA embryos (**Figure 3D and S3C**). As *Ptbp1* and *Srsf10* are downregulated in Val_LNA embryos (**Figure 3A**), *Srsf10* transcripts with AL events and *Ptbp1* transcripts with SE events are potentially unstable and degraded by non-sense-mediated decay pathways. *Rbm39*, on the other hand, was upregulated in Val_LNA embryos, and its transcripts showed less skipping of exon 2b in Val_LNA embryos than Ctrl_LNA embryos (**Figure 3E-F and S3E**). We also detected events with alternative 3’ splice sites (A3). The top gene with A3 was *Mus81*, a gene known to initiate compensatory DNA synthesis during mitosis and resolve mitotic interlinks, thereby facilitating proper chromosome segregation [56–58]. While *Mus81* was not differentially expressed in Val_LNA embryos relative to Ctrl_LNA embryos (**Figure 3G**), *Mus81* transcripts showed lower levels of inclusion of an alternative 3’ splice site in Val_LNA embryos (**Figure 3H, S3F**). These data demonstrated that splicing was affected in Val_LNA embryos, including alternative splicing of splicing factors.

### tRFValCAC regulates alternative splicing of Pyruvate kinase gene

We further explored the role of tRFValCAC in regulating the splicing of embryo-specific isoforms. Pyruvate kinase gene *Pkm* has two isoforms, *Pkm1* and *Pkm2* [59, 60] (**Figure 4A**). PKM2 is an embryonic isoform that is also known to be expressed in cancer cells and promotes aerobic glycolysis [61, 62]. PKM1 is the adult isoform that promotes oxidative phosphorylation [61]. These two isoforms result from mutually exclusive alternative splicing of the *Pkm* pre-mRNA, including exon 9 (*Pkm1*) or exon 10 (*Pkm2*). hnRNPA1, another member of the hnRNP family of proteins and a splicing regulator, has been previously shown to promote the inclusion of exon 10 and the generation of the *Pkm2 isoform* [59, 63]. hnRNPA1 can bind repressively to sequences flanking exon 9, resulting in exon 10 inclusion. RNA-binding motif protein X-linked (RBMX, also known as hnRNPG) modulates hnRNPA1-mediated alternative splicing of *Pkm* by inhibiting hnRNPA1 binding to sequences flanking *Pkm* exon 9, leading to lower *Pkm2* levels. [64].

**Figure 4:**
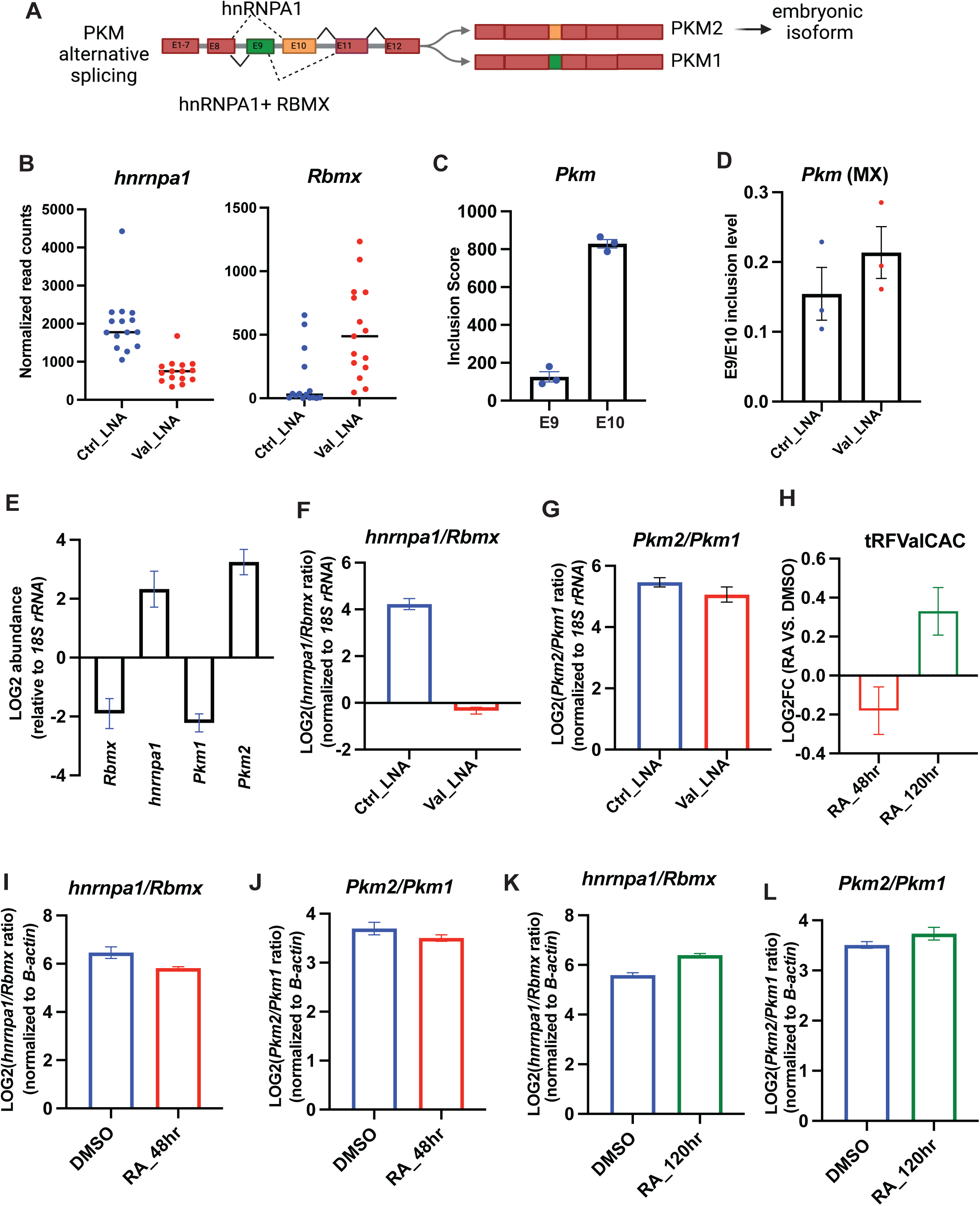
tRFValCAC regulates alternative splicing of *Pkm* isoforms. **A)** Schematic showing alternatively spliced forms of *the Pkm* gene and the role of hnRNPA1 and RBMX in regulating splicing of *Pkm1* and *Pkm2* isoforms. **B)** Single embryo mRNA-seq normalized read counts of *hnrnpa1* and *Rbmx* transcripts (DESeq2 *padj* values: *hnrnpa1* 2.73×10^-9^ and *Rbmx* 0.10). **C)** Alternative splicing analysis of *Pkm* transcripts in 2-cell control embryos (Ctrl_LNA embryos). The bar graph shows the inclusion score for exon 9 and exon 10 in *Pkm* transcripts. Consistent with *Pkm2* being an embryo-specific isoform, exon 10 is included at a significantly higher level compared to exon 9. **D)** Analysis of alternative splicing of *Pkm* transcripts in Val_LNA embryos compared to Ctrl_LNA embryos. Val_LNA embryos showed a higher exon 9 (E9) /exon 10 (E10) inclusion ratio, indicating an increase in non-embryonic *Pkm1* isoform in Val_LNA embryos. **E)** Single embryo qRT-PCR was performed to determine the abundance of specific transcripts in control embryos (Ctrl_LNA). Data is represented as log2 fold change relative to 18S rRNA. **F)** Single embryo qRT-PCR data showing *hnrnpa1/Rbmx* ratio in Val_LNA compared to Ctrl_embryos (normalized to 18S rRNA). There was a significant decrease in *hnrnpa1/Rbmx* ratio in Val_LNA embryos (unpaired t-test; p-value < 0.0001. **G)** Single embryo qRT-PCR to determine *Pkm2/Pkm1* ratio using isoform-specific primers (see Methods). There was a subtle decrease in *Pkm2/Pkm1* ratio in Val_LNA embryos compared to Ctrl_LNA embryos. 4 Ctrl_LNA and 5 Val_LNA single embryos were analyzed by qRT-PCRs in (F) and (G). **H)** Log2 fold change in abundance of tRFValCAC in mESCs treated with RA for 48 hours and 120 hours relative to DMSO treatment (normalized to *B-actin*; n=3 replicates). **I)** qRT-PCR in mESCs treated with RA for 48 hours, which leads to a reduction in tRFValCAC levels (as shown in (H)). The bar graph shows *the hnrnpa1/Rbmx* ratio in DMSO and RA-treated mESCs (normalized to B-actin). *hnrnpa1/Rbmx* ratio is significantly reduced in RA treated cells (n= 9 replicates, unpaired t-test *p* value= 0.0196). **J)** qRT-PCR to determine *the Pkm2/Pkm1* ratio in mESCs treated with DMSO (n=8) or RA (n=9) for 48 hours. **K)** qRT-PCR in mESCs treated with RA for 120 hours to induce tRFValCAC (as shown in (H)). The bar graph shows *the hnrnpa1/Rbmx* ratio in DMSO and RA-treated mESCs (normalized to *B-actin*). *hnrnpa1/Rbmx* ratio is significantly increased in RA treated cells (n= 6 replicates, unpaired t-test *p* value <0.0001). **L)** qRT-PCR analysis showing *Pkm2/Pkm1* ratio in DMSO and RA treated mESCs (normalized to *B-actin*) (n=8 (DMSO) and 7 (RA)). Graphs represent data as mean ± SEM.

We noticed that Val_LNA embryos displayed a decrease in *hnrnpa1* transcript abundance and an increase in *Rbmx* levels compared to Ctrl_LNA embryos (**Figure 4B**), revealing an overall decrease in *hnrnpa1/Rbmx* ratio upon inhibition of tRFValCAC. Therefore, we next investigated whether inhibition of tRFValCAC leads to alterations in *Pkm1* and *Pkm2* isoform levels.

Consistent with *Pkm2* being the embryonic isoform, our alternative splicing analysis revealed that embryonic *Pkm* transcripts predominantly include exon 10 (**Figure 4C**). Interestingly, we detected a slight increase in exon 9 inclusion in Val_LNA embryos compared to Ctrl_LNA embryos (**Figure 4D).** To further analyze changes in *Pkm* splicing in Val_LNA embryos, we examined the abundance of specific *Pkm* isoforms using single embryo qRT-PCRs and isoform-specific primers. Single-embryo qRT-PCRs on control embryos also showed that *hnrnpa1* is more abundant than *Rbmx* and confirmed that *Pkm2* is the major embryonic isoform (**Figure 4E**). Consistent with mRNA-seq analysis, single embryo qRT-PCR revealed a significant decrease in *hnrnpa1/Rbmx* transcript ratio in Val_LNA embryos compared to Ctrl_LNA embryos (**Figure 4F**). Furthermore, isoform-specific qRT-PCR revealed a slight decrease in *Pkm2/Pkm1* ratio in Val_LNA embryos, consistent with mRNA-seq data (**Figure 4G**).

Next, we turned to mESCs to further examine the specificity of the effects of tRFValCAC on *Pkm* splicing using RA treatment. As reported previously [55], we noted an initial downregulation of tRFValCAC levels after 48 hours of RA treatment, followed by an increase after 120 hours of RA treatment in mESCs (**Figure 4H**). Thus, the timing of RA treatment allowed us to bidirectionally alter endogenous levels of tRFValCAC in mESCs and examine whether *hnrnpa1*/*Rbmx* ratio correlated with changes in tRFValCAC levels. Indeed, reduced tRFValCAC levels in response to 48hrs RA treatment led to a decrease in *hnrnpa1*/*Rbmx1* ratio and a slight reduction in *Pkm2/Pkm1* ratio (**Figure 4I-J**), consistent with changes observed in Val_LNA embryos. Moreover, increased tRFValCAC levels in response to 120 hours of RA treatment led to increased *hnrnpa1*/*Rbmx* and *Pkm2/Pkm1* ratios (**Figure 4K-L**). Collectively, our findings from both embryos and mESCs indicate that tRFValCAC regulates the levels of *hnrnpa1* and *Rbmx* transcripts and, consequently, positively influences the levels of the embryonic *Pkm2* isoform and potentially embryonic metabolism.

To delve deeper into the potential mechanism of tRFValCAC-mediated regulation of gene expression and RNA splicing in embryos, we profiled interacting proteins of tRFValCAC in mESCs (**Table S5**). Intriguingly, the top proteins that showed significantly higher abundance in tRFValCAC-pulldown relative to the scrambled control pulldown were ribosomal proteins and hnRNPs, including hnRNPA2B1, hnRNPA1, hnRNPAB and hnRNPM. GO analysis also showed that mRNA processing, post-transcriptional regulation of gene expression, translation, and RNA splicing were among the top enriched pathways (**Table S6**). Notably, the splicing regulator hnRNPA1 was identified as a potential interacting protein, suggesting that the regulation of *hnrnpa1/Rbmx* ratio and *Pkm* splicing may involve direct binding of tRFValCAC to hnRNPA1. Overall, tRFValCAC-interacting proteins identified here suggest that tRFValCAC regulates gene expression by interacting with RNA-binding proteins, including splicing factors and ribosomal proteins.

### tRFValCAC regulates preimplantation embryonic development

Given the dramatic alterations in transcript abundance in 2-cell embryos upon inhibition of tRFValCAC, we next investigated the role of tRFValCAC in regulating preimplantation embryonic development. We introduced Val_LNA or Ctrl_LNA into embryos and monitored the dynamics of the first cleavage. We observed abnormalities during the initial cell cleavage in the Val_LNA injected embryo. Specifically, numerous Val_LNA embryos either did not complete the first cleavage or were stuck at that stage (**Figure 5A**). Next, we examined the effects of tRFValCAC inhibition on preimplantation embryonic development until the blastocysts stage. We again observed that ∼20% of Val_LNA embryos did not progress beyond the 2-cell stage even 48 hrs post IVF (**Figure 5B**). Beyond 48 hours, Val_LNA embryos showed further delayed development; while 88% of Ctrl_LNA embryos reached the blastocyst stage, only 58% of Val_LNA embryos progressed to the blastocyst stage, with the remaining lagging at the morula stage (**Figure 5C-D**). These data suggest that tRFValCAC-mediated gene expression regulation in 2-cell embryos can modulate preimplantation embryonic development.

**Figure 5:**
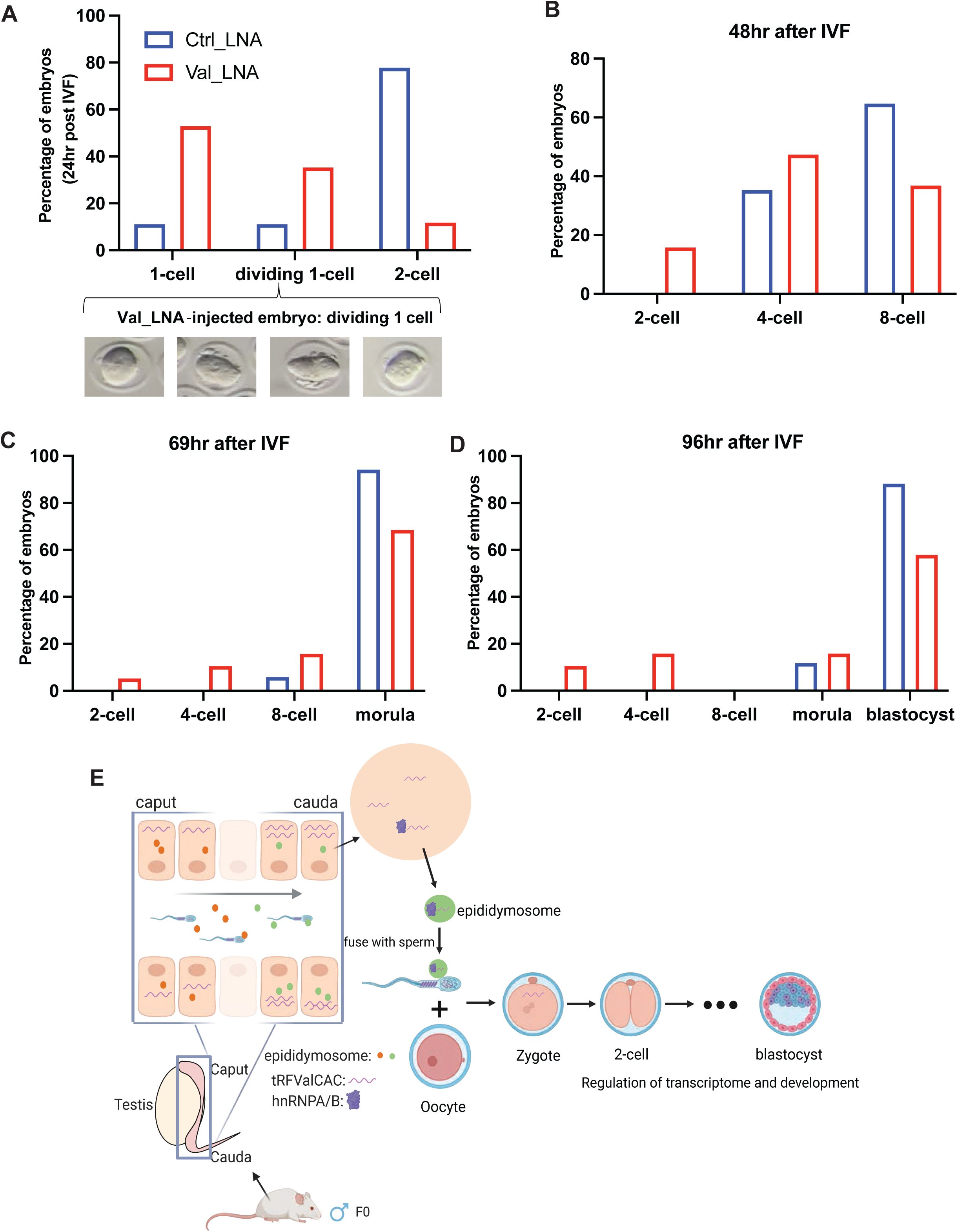
Val_LNA embryos display delayed preimplantation development. Post-IVF, the male pronucleus was injected with Val_LNA or Ctrl_LNA, and the embryos were cultured in-vitro until the blastocyst stage to monitor embryonic development upon inhibition of tRFValCAC. **A)** 1-cell and 2-cell stage embryos as the proportion of the total embryos injected (excluding dead and amorphous embryos). A subpopulation of embryos was stuck at the first cell division. The images below the graph are representative images of such embryos. Ctrl_LNA n=11, Val_LNA n=7 **B-D)** IVF-generated embryos were injected with Ctrl_LNA (n=17) or Val_LNA (n=19), and their developmental stages were examined over a period of 96 hrs. The percentage of embryos at a specific development stage was calculated as proportion of total embryos analyzed at a given time point. **E)** A schematic of the proposed mechanism of tRFValCAC dynamics in sperm and its regulatory role in embryonic development. Our data support a model wherein tRFValCAC, in complex with hnRNPAB, is delivered from epididymis to sperm via epididymosomes. tRFValCAC is then deposited by sperm into the oocyte upon fertilization (likely still in complex with hnRNPAB), where it regulates early embryonic gene expression, RNA processing, and development.

## DISCUSSION

Here, we identified the potential role of an RNA-binding protein hnRNPAB in regulating sperm tRFValCAC levels and uncovered a novel function of tRFValCAC in regulating early embryonic transcriptome and development (**Figure 5E**).

We previously reported that tRFValCAC is gained by sperm during post-testicular maturation in the epididymis and that it can be delivered to sperm from the epididymis via epididymosomes secreted from epididymis epithelial cells [4, 5]. Here, we set out to investigate the mechanistic basis of tRFValCAC delivery from epididymis to sperm by focusing on proteins that interact with tRFValCAC. We found that an RNA-binding protein, hnRNPAB, binds directly to tRFValCAC in the epididymis and that SUMOylated form of hnRNPAB is the most prominent form in the epididymis and epididymosomes. Interestingly, we found that the knockdown of hnRNPAB in epididymis epithelial cells led to reduced levels of tRFValCAC in the EVs secreted from those cells, suggesting a role of hnRNPAB in sorting tRFValCAC into EVs. Our data support a model wherein tRFValCAC binds to hnRNPAB in the epididymis, and this binding facilitates its sorting into epididymosomes and, thereby, its delivery to sperm. Other members of the hnRNP family have been reported to regulate miRNA sorting into EVs, including hnRNPA2B1 [43, 65], hnRNPC1 [66], and SYNCRIP (also known as hnRNPQ) [67], with some interactions being dependent on specific sequence motifs in miRNAs and SUMOylation of the hnRNP protein. Pull down of hnRNPAB interacting small RNAs in the epididymis revealed that tRFValCAC specifically binds to hnRNPAB (**Figure S1D**) (no other tRFs were detected in the pull-down, except for a fragment of tRNA-GlyGCC). Therefore, this interaction is likely based on the unique sequence, secondary structure, or modifications of tRFValCAC. The SUMOylated form of hnRNPAB does not seem to regulate the sorting of tRFValCAC into EVs as only the non-SUMOylated form of hnRNPAB was detected as an interacting partner of tRFValCAC in DC2 cells (wherein its knockdown led to reduced levels in the EVs).

Functionally, we uncovered that inhibition of tRFValCAC in preimplantation embryos altered transcript abundance of many genes, including those involved in cell cycle, chromosome organization, and RNA splicing. Moreover, these embryos showed alternative splicing events, including alternative splicing of splicing regulators *Srsf10*, *Rbm39*, and *Ptbp1*. These changes in splicing could be consequence of changes in the abundance of splicing regulators. Several cases of splicing factors autoregulating their splicing have been reported, including RBM39 [68], SRSF10 [69], and PTB [70]. For instance, here we found that *Srsf10* was downregulated in Val_LNA embryos, and there was an increase in the inclusion of an alternate last exon (AL) in the *Srsf10* transcript. Including an alternate last exon could result in a transcript degraded by the nonsense-mediated decay pathway [71], reducing the overall transcript abundance of *Srsf10* [69]. Focusing further on specific misregulated genes in Val_LNA embryos, we found that the abundance of *hnrnpa1* and *Rbmx* transcripts was altered. hnRNPA1 and RBMX are known to regulate the splicing of the *Pkm* gene. We found that, both in embryos and mESCs, loss-of-function and gain-of-function of tRFValCAC resulted in subtle downregulation and upregulation of Pkm2/Pkm1 isoform ratio, respectively. As *the Pkm2* isoform is critical for proliferation and growth [72], a slight decrease in the Pkm2/Pkm1 ratio in the Val_LNA embryos could affect embryonic development. Moreover, alterations in transcript abundance of genes involved in cell-cycle regulation and chromatin organization can also modulate early embryonic development. Consistently, Val_LNA embryos had delayed development compared to control embryos.

What is the mechanism of tRFValCAC-mediated regulation of embryonic transcriptome? Based on known mechanisms of gene regulation by tRFs, tRFValCAC could regulate transcript abundance at the level of transcription [18, 19, 25, 73], mRNA stability [15, 24], or translation [10, 11, 14]. We found that in mESCs, tRFValCAC interacts with various proteins involved in gene regulation, notably those from the hnRNP family and ribosomal proteins. Binding to ribosomal proteins can directly influence translation. Indeed, a prior study reported that a 5’ fragment of tRNA-Valine-CAC binds to ribosomes and inhibits global translation during stress [14]. hnRNPs are a large class of well-characterized RNA-binding proteins involved in regulating mRNA transcription, splicing, editing, translation, stability, and localization [46, 74, 75]. Thus, tRFValCAC potentially regulates early embryonic gene expression by interacting with these RNA-binding proteins and the transcript abundance changes observed upon inhibition of tRFValCAC are likely a result of alteration of mRNA processing or translational efficiency of upstream regulators. We previously found that a 5’ fragment of tRNA-Glycine-GCC (tRFGlyGCC), another tRF highly abundant in mature sperm, regulates transcription of a specific set of retroviral element-driven genes in preimplantation embryos [26]. A tRF derived from tRNA-Gln-TTG (tRFGlnTTG) was also shown to regulate retroviral elements and cell-cycle genes in pig embryos [76]. It was recently reported that tRFGlyGCC regulates its target genes in mESCs by regulating the biogenesis of other small non-coding RNAs, specifically U7 snRNA, which in turn regulates histone pre-mRNA processing and, consequently, chromatin organization [26]. Furthermore, tRFGlyGCC was shown to interact with two related members of the hnRNP family, hnRNPF and hnRNPH. Given that we also isolated hnRNP proteins as potential interacting partners of tRFValCAC in mESCs, these observations suggest that hnRNP proteins are likely effectors of tRF functions in early embryos.

Finally, we discuss the implications of these findings for small RNA-mediated intergenerational epigenetic inheritance of paternal environmental effects. There is a growing body of work demonstrating that levels of sperm tRFs are altered in response to various environmental perturbations in mice [4, 30, 31, 33, 34, 37, 77–79] and humans [39, 80], and injection of tRFs isolated from sperm of exposed mice into control zygotes can lead to metabolic disruption in the resulting offspring [30, 33, 79]. How tRFs regulate offspring metabolic phenotypes remains unclear. Here, we found that one of the highly abundant tRFs in sperm, tRFValCAC, regulates early embryonic gene expression, and inhibition of tRFValCAC leads to delayed preimplantation development. Paternal dietary challenges have been shown to affect preimplantation development. For instance, we previously found that paternal low protein lowers the number of blastocyst cells [4]. Paternal high-fat diet-induced obesity was also reported to alter the pace of preimplantation embryonic development, with decreased cleavage and reduced development to the blastocyst stage [81, 82]. Moreover, defective preimplantation development can lead to metabolic diseases in adulthood [83, 84]. Thus, our results support a model wherein the paternal environment can alter the levels of sperm deposited tRFs in the embryo, which in turn can modulate early embryonic gene expression and development and, consequently, metabolic phenotypes in adulthood. Taken together, our study sheds light on the potential mechanism of sperm tRF-mediated intergenerational epigenetic inheritance.

## METHODS

### Mice and cell lines

Wild-type FVB/NJ mice were used in the study. All animal care and use procedures were in accordance with the guidelines of the University of California Santa Cruz Institutional Animal Care and Use Committee. Mice were group-housed (maximum of 5 per cage) with a 12-hour light-dark cycle (lights off at 6 pm) and free access to food and water *ad libitum*. E14 mouse embryonic stem cell (mESCs) were cultured in high-glucose DMEM (Gibco, cat# 11965118), 10% FBS (Gibco, cat# A5256701), 1% non-essential amino acids (Gibco, cat# 11140076), 1% glutamine (Gibco, cat# 35050061), 0.1mM beta-mercaptoethanol (Sigma, cat# M6250), and LIF (Sigma, cat# ESG1107). For tRFValCAC induction, mESCs were treated with 0.5 μM Retinoic acid (RA) or DMSO for 48 hr or 120 hr, according to a previous report [55]. Distal caput epididymis-derived immortalized cell line (DC2) was a gift from Marie-Claire Orgebin-Crist. DC2 cells were seeded in 75 cm^2^ Nunc EasYFlasks (Thermo Fisher, cat # 156499) coated in collagen type 1, rat tail (Gibco, cat # A1048301). Cells were cultured in Iscove’s modified Dulbecco’s medium (IMDM) supplemented with 10% exosome-free fetal bovine serum (Gibco, cat# A2720801), 1% penicillin-streptomycin (Gibco, cat# 15140122) and 0.3ul/ml DHT (Millipore, cat# 521-18-6).

### Preparation of single-cell suspension of epididymides

Epididymides were dissected and made into single-cell suspension following a published protocol with modifications [85]. Epididymides from a 10- to 12-week-old FVB mouse were dissected and sliced into 1–5 mm fragments, then incubated in prewarmed 1X PBS. Once the tissues were cleared of any visible sperm, they were transferred to a glass flask containing freshly prepared tissue dissociation media (DMEM, Collagenase IV 4 mg/ml, DNAse I 0.05 mg/ml) and placed on a shaker rotating at 200 rpm at 35°C for 30 to 45 minutes. Next, samples were allowed to settle, and 5 ml of supernatant was removed. 10 ml 0.25% trypsin-EDTA and 0.05 mg/mL DNAse I solution were added to the flask, and the flask was shaken for another 50 minutes until there were no observable tissue pieces. The resulting solution was sequentially filtered through 100 μm, 70 μm, and 40 μm cell strainers, followed by centrifugation and washing of the pellet with 1XPBS. The cell pellets were subjected to lysing using cell lysis buffer (20 mM Tris–HCl pH 7.5, 150 mM NaCl, 1.8 mM MgCl2, 0.5% NP40, protease inhibitor cocktail for mammalian cells (no EDTA), 1 mM DTT, 80 U/ml RNase inhibitor). The resulting epididymal cellular extract was used as the input for the pull-down of tRFValCAC-interacting proteins using a biotinylated RNA oligo, as described below.

### tRFValCAC-interacting proteins pull-down

Protein pulldowns with 3’-biotinylated RNA mimics were conducted using a previously published protocol [55]. Briefly, 1ug of biotinylated tRFValCAC synthetic oligo or a scrambled control oligo in 250 ul of binding buffer (10 mM Tris pH 7.0, 0.1 M KCl, 10 mM MgCl2) was incubated with 500 ug of epididymal cellular extract. Next, RNA-bound proteins were eluted in Laemmli buffer using streptavidin agarose beads. 2× Laemmli buffer was added to the protein samples, boiled at 95°C for 10 minutes, and loaded on an SDS-PAGE gel. The gel was run for a few minutes to allow proteins to migrate out of the wells when protein-containing gel pieces were cut out and submitted for mass spectrometry analysis at the Vincent J. Coates Proteomics/Mass Spectrometry Laboratory at the University of California, Berkeley. A similar procedure was used to validate protein interaction through Western blot, but the gel was allowed to run until the dye front reached the end of the gel. The gel was transferred onto a PVDF membrane and subsequently probed with antibodies against hnRNPAB (Santa Cruz Biotechnology, cat # sc-376411).

### LC-MS/MS data analysis

The EmPAI [86] method was used to analyze the MS data from tRFValCAC-interacting proteins pulldown from the epididymis. PAI (protein abundance index) represents the number of peptides per protein normalized by protein size. PAI was then converted to exponentially modified PAI (emPAI), equal to 10PAI minus one, proportional to protein content in a protein mixture. Zero EmPAI (protein abundance index) values were substituted with the lowest detected values (0.01). Then, scramble control emPAI values were subtracted from those of tRFValCAC pulldown. Any candidates exhibiting higher emPAI scores in scramble controls compared to the tRFValCAC pulldown were excluded, and the tRFValCAC-interacting proteins ranked with high emPAI to low. Proteins were considered significantly enriched in the sample at an FDR-adjusted p-value < 0.05. Identification of tRFValCAC-interacting proteins in mESCs was performed the same way described above for the epididymis. The mESC pulldown LC-MS data was analyzed using PEAKS Studio 11 software. Briefly, raw MS data were processed through PEAKS Studio 11 for high-confidence identification of proteins. Proteins that did not pass the filter for significant peptides were removed from further analysis. Next, spectral counts were used to compare protein abundance in tRFValCAC pulldown and scramble control samples. Proteins that showed higher spectral counts in the control compared to the tRFValCAC pulldown were excluded from further analysis. The filtered protein list was then analyzed for significantly higher abundance in tRFValCAC pulldown using a t-test with a p-value cutoff of <0.05.

### Epididymis, sperm, and epididymosome collection

Epididymides and sperm were collected from male mice (10-12 weeks old) as previously described [4]. Caput and cauda epididymides were dissected out and transferred to a 35mm dish containing 1ml of pre-warmed Whitten’s Media (100 mM NaCl, 4.7 mM KCl, 1.2 mM KH2PO4, 1.2 mM MgSO4, 5.5 mM Glucose, 1 mM Pyruvic acid, 4.8 mM Lactic acid (hemicalcium), and HEPES 20 mM). Two incisions were made in the tissue, and the tissue was gently squeezed to release fluid. After a 15-minute incubation at 37°C, media were transferred to a new tube for an additional 15-minute incubation, and epididymis tissues were flash-frozen in liquid nitrogen. After incubating for 30 minutes at 37 °C, the media was spun at 2000 x g for 2 minutes. The supernatant that contained epididymosomes was transferred to a fresh 1.5 ml tube, and the sperm pellet was washed with 1X PBS, followed by incubation in lysis buffer (0.1% SDS and 0.5% Triton-X) for 10 minutes on ice to remove somatic cell contamination. The sperm sample was finally washed with 1X PBS and pelleted after centrifuged at 2000 x g. The supernatant with epididymosome was centrifuged at 10000 x g for 30 minutes to remove cellular debris, followed by ultracentrifugation at 120,000 × g at 4 °C for 2 hours (Beckman, TLA120.4 rotor). The resulting pellets were washed in cold PBS and underwent a second ultracentrifugation at 120,000 × g at 4 °C for 2 hours. These epididymosome pellets were then resuspended in RIPA lysis solution for protein extraction.

### Western blot analysis

Protein extracts were denatured in 2X Laemmli buffer (Bio-Rad, cat# 1610737) at 95°C for 5 minutes and subjected to electrophoresis on a 4-20% SDS-PAGE gel. Subsequently, proteins were transferred onto a PVDF membrane. The membrane was then incubated with anti-hnRNPAB (Santa Cruz Biotechnology, cat # sc-376411), SUMO1 Monoclonal Antibody (Invitrogen, cat # 21C7), or Anti-beta Actin (Abcam, cat # ab8227) antibodies at 1:1000 dilution followed by secondary antibody treatment at 1:10000 dilution (Goat anti-Mouse, cat # 31430, Invitrogen).

### UV-RNA immunoprecipitation assay

The immunoprecipitation was conducted following the methods outlined in a previous study [55]. Cells with UV-crosslinked at 450 mJ/cm2 and following crosslinking cell extracts were prepared using a lysis buffer containing 100 mM Tris pH 7.4, 150 mM NaCl, 1% NP40, RNase inhibitor, protease inhibitor cocktail, and 1 mM DTT. The cell extract was then pre-cleared with protein A/G resin (Thermo Scientific, cat # 53132) for 1 hour at 4°C, followed by incubation with either hnRNPAB antibody (Santa Cruz Biotechnology, cat # sc-376411) or IgG control (Cell Signaling Technologies, cat # 3900S) overnight at 4°C. After four washes with lysis buffer, RNA from the protein complex was released by proteinase K treatment at 37°C for 10 minutes. The RNA was subsequently extracted using TRIzol (Invitrogen, cat # 15596026). Small RNA libraries were constructed using the OTTR-seq method, described below [87].

### DC2 cell EV isolation

DC2 cell culturing and vesicle isolation were processed as previously described [88]. EVs were isolated from DC2 cell-culture media using sequential ultracentrifugation. Briefly, cellular debris was removed from the media by centrifugation at 200 × g for 10 minutes, 2000 × g for 10 minutes, and 10,000 × g for 30 minutes. Next, EVs were pelleted by ultracentrifugation at 55,000 × rpm for 2 hours using the TLA120.4 rotor (Beckman). The EV pellet was resuspended in 1XPBS and frozen at −80 °C until further analysis.

### Quantitative real-time PCR

RNA extraction followed the method described above. Each sample utilized a minimum of two biological replicates. Following the manufacturer’s protocol, reverse transcription (RT) was performed using the SuperScript III Kit (Invitrogen, cat # 18080093). Quantitative real-time PCR (qRT-PCR) was carried out using the KAPA SYBR FAST qPCR mix (KAPA Biosystems). The primer details for qRT–PCRs can be found in **Table S7.**

### TaqMan Small RNA assays

tRFValCAC and U6 levels were examined using custom-designed TaqMan small RNA Assays, following the manufacturer’s protocol (ThermoFisher). Briefly, 10 ng of RNA was reverse transcribed using the TaqMan reverse transcription kit (ThermoFisher, cat # 43-665-96).

Following this, qRT-PCR was performed in 15 µL reactions employing TaqMan Universal PCR Master Mix (ThermoFisher, cat # 43-643-38), according to the manufacturer’s PCR program.

### Small RNA sequencing

OTTR-seq small RNA sequencing was performed as previously described [87]. Briefly, input RNA was labeled at the 3’ end by incubation in a buffer containing ddATP for 90 minutes at 30°C, followed by the addition of ddGTP and another incubation at 30°C for 30 minutes. The reaction was stopped by incubating at 65°C for 5 minutes, followed by adding 5 mM MgCl2 and 0.5 units of shrimp alkaline phosphatase (NEB, cat # M0371S) at 37°C for 15 minutes. The reaction was stopped by adding 5 mM EGTA and incubating at 65°C for 5 minutes. Samples were then incubated in templated cDNA synthesis buffer, adaptors, and dNTPs at 37°C for 20 minutes, followed by heat inactivation at 65°C for 5 minutes and RNase A/H treatment. cDNA was size selected on 10% PAGE-Urea gel to minimize adaptor dimers. Size selected cDNA was PCR amplified for 12 cycles with Q5 high fidelity polymerase (NEB, cat # M0491S). The final PCR product was cleaned with AMPure XP beads (Beckman, cat # A63881) and separated by 6% PAGE gel to remove adaptor dimers. The desired product was excised from the gel and eluted in 400ul elution buffer overnight at -20°C, followed by isopropanol precipitation and resuspension in 9µl of RNase-free H2O. The final libraries were sequenced on a HiSeq or NextSeq instrument.

Small RNA sequencing data analysis was done using the tRAX analytical analysis tool [89] with default parameters except for a minimum non-tRNA read size of 16 and using the Ensembl gene set [90], the mm10 piRNA gene set from piPipes [91], and ribosomal RNA repeats from UCSC repeatmasker [92] as the gene set. In tRAX, reads were mapped to the mouse mm10 genome combined with tRNA sequences taken from gtRNAdb [93]. tRNA reads were defined as any reads whose best mapping includes a mature tRNA sequence. The tRAX pipeline uses bowtie2 with options “-k 100 --very-sensitive --ignore-quals --np 5 –very-sensitive,” extracts all best mappings from those results, and categorizes all tRNA mappings to acceptor type-specific, decoder type-specific, or unique tRNA transcript-specific, and only reads specific to acceptor type and anticodon were used for corresponding tRNA counts. Reads that mapped to mature tRNAs were further classified into four fragment types based on the read alignment and the ends of the tRNAs. Reads where the 5’ end lies within 3 nts and 3’ end lies with 5 nts of the respective ends on the mature tRNAs are categorized as “whole” full-length tRNAs (referred to as “tRNAs”), and reads that overlap or closely align to either the 5’ ends or 3’ end of the tRNAs are classified as “tRF_5’” and “tRF_3’”, respectively. Mature tRNA reads that do not display these features are classified as “tRF_other”. Adjusted p-values and log2-fold change were calculated using DESeq2 [94] with default parameters as a component of the tRAX pipeline, and plots were generated with ggplot2 and Prism.

### Transfection of mESC and DC2 cells

Cell transfections were conducted in OptiMEM using 6-well plates [4]. Briefly, mESCs were transfected with 1ug of psiCHECK-2 plasmid with either 1ng of Val_LNA oligo or Ctrl_LNA oligo using Lipofectamine 2000 (Invitrogen, cat # 11668019). DC2 cells were transfected with 400 ng of siRNA targeting hnRNPAB using Lipofectamine 2000. After 16 hours, media was changed, and cells were allowed to grow for a total of 48 hours when RNA extraction was performed using the standard Trizol protocol.

### Dual reporter assay in mESC

The Dual-Luciferase® Reporter Assay (Promega, cat # E1910) was conducted following the manufacturer’s instructions. tRFValCAC sequence was inserted into the multiple-cloning site at the 3’ end of the *hRluc* gene within the psiCHECK-2 Vector (Promega, cat # C8021). mESCs were transfected using the method described above, wherein LNA_Val or Ctrl_LNA oligo was co-transfected with the tRFValCAC containing psiCHECK2 vector. The transfected mESCs were then employed for the Dual-Luciferase® Reporter Assay to evaluate Renilla luciferase activity, while Firefly luciferase activity was utilized as an internal control for plasmid expression.

### Northern blot analysis

Northern blot was performed following the previous protocol [95]. Briefly, total RNA from mESC was mixed with an equal amount of Gel loading buffer II (Invitrogen, cat # AM8546G) and heated to 65°C for 5 minutes to denature the RNA. The RNA was then run on a 10% PAGE 7M urea denaturing gel, followed by transfer to a positively charged nylon membrane and then crosslinked with EDC (1-Ethyl-3-(3-dimethylaminopropyl)carbodiimide) at 60°C for 1–2 hours and prehybridized with ULTRAhyb Ultrasensitive Hybridization Buffer (Invitrogen, cat # AM8670). The membrane was then hybridized with 50 pmol mL−1 Biotin-labeled Locked Nucleic Acid-modified DNA probes (designed and synthesized by Qiagen) specific to the 5’ end of tRNA-ValCAC and 5S rRNA. After washing, the blots were then processed and developed using a Chemiluminescent Nucleic Acid Detection Module Kit (ThermoFisher Scientific, cat # 89880). The Sequences of oligos used are: tRNA-ValCAC 5’ CGT GAT AAC CAC TAC ACT ACG G /3Bio/ 3’ and 5’ GAT CGG GCG CGT TCA GGG TGG TAT G/3Bio/ 3’.

### In vitro fertilization (IVF)

The protocol was performed following a previously reported method with some modifications [4]. FVB/NJ mice were used as egg and sperm donors. Cauda epididymis and vas deferens sperm were collected from 8-12 weeks old males. The tissue was placed in 1ml prewarmed EmbryoMax® Human Tubal Fluid (HTF) (Sigma, cat # MR-070-D) media containing 0.75mM Methyl-beta-cyclodextrin, and sperm was allowed to swim out by incubating the tissue at 37°C for 20-30 minutes. Sperm were counted, and 1×10^6^ sperm was then pre-incubated for 1 hour at 37°C and 5% CO2 in 200 μl drops of fertilization medium KSOM (EmbryoMax® Advanced KSOM Embryo Medium, Sigma, cat # MR-101-D) + 1mM Glutathione covered with sterile mineral oil. Superovulation was induced in female mice using PMSG and hCG as in [4].

Cumulus-oocyte complexes were collected from the oviducts of females at 13–15 h after hCG injection. These were then transferred to the 200uL fertilization drop. The oocytes were then co-cultured with sperm at 37°C under 5% CO2 for 4 hours to allow fertilization. Fertilized zygotes were washed to get rid of excess sperm and cumulus cells and cultured to later stages of development at 37°C under 5% CO2 and 5% O2.

### Embryo RNA microinjection experiments

Protocol was adopted from [4]. Zygotes for microinjection studies were generated by IVF as described above. After four hours of IVF, the zygotes were washed three times in KSOM medium and placed in a drop of KSOM medium at 37°C in 5% CO_2_ and 5% O_2_ for 2 hours. Embryos were then transferred to a 10ul KSOM drop covered by oil and subjected to micromanipulation. Embryos were microinjected with either H3.3-GFP mRNA with LNA-containing scramble oligo (Ctrl_LNA) (control group) or H3.3-GFP mRNA plus tRFValCAC antisense LNA oligo (Val_LNA) (experimental group). RNA injections were carried out using a Femtojet (Eppendorf) microinjector at 100 hPa pressure for 0.2 seconds, with 7 hPa compensation pressure. RNAs used for microinjections and their concentrations were 100 ng/µl of H3.3-GFP mRNA, 200 ng/µl of control oligo (5’ AACACGTCTATACGC 3’), and 200 ng/µl of 5’tRFValCAC-2 antisense LNA oligo (5’ CGAACGTGATAACC 3’). After the microinjections, embryos were placed back into the incubator, and H3.3-GFP fluorescence was verified at the 2-cell stage. GFP-positive embryos were cultured until the late 2-cell stage for mRNA-seq (28 hours post-IVF) or until the blastocyst stage for developmental analysis.

It should be noted that as 5’ sequence of tRNA-Val-CAC-2, tRNA-Val-CAC-4, and tRNA-Val-AAC-1 are highly similar, the 5’ fragments derived from these tRNAs are indistinguishable for the most part and, therefore, this LNA oligo is predicted to also target 5’ fragments derived from tRNA-Val-CAC-4 and tRNA-Val-AAC-1. Given that the 5’ fragments generated from these tRNAs will be of identical sequence, it is likely that they are a single entity in the cell.

Alternatively, the 5’tRFs derived from these three tRNAs could differ in their RNA modification status and have distinct functions.

### Single embryo mRNA-Seq

Single embryo mRNA-Seq libraries were generated using the SMART-Seq protocol [96] as described previously [4]. Embryos with fewer than 10,000 sequenced reads were excluded from the dataset. The 151x151bp paired-end sequencing reads were trimmed at the 3’ end to 120x120bp. The trimmed reads that mapped to the mouse genome repeat elements (mostly rRNA) were removed from further processing. The repeat-filtered reads were aligned to the mouse (mm10) genome (augmented by a splice junction database derived from GENCODE transcriptome version VM18) with STAR, version 2.5.3a, with soft-clipping disabled (using the parameter alignEndsType EndToEnd). Duplicate alignments were removed, leaving about 6.5 million mapped reads per sample. Differential gene expression from these mapped reads was assessed using DESeq2, version 1.38.3, comparing 15 replicates each of the “Val_LNA” (experimental group) and “Ctrl_LNA” (control group) conditions. The GO analyses for significantly altered genes were done using PANTHER and Broad Institute’s GSEA tools.

### Alternative splicing analysis of RNA-seq data

For splicing analysis, the mapped reads were combined into 3 pseudo-replicates for each of the “Val_LNA” and “Ctrl_LNA” conditions, comprising 5 or 4 replicates per pseudo-replicate, chosen to maximize the difference between pseudo-replicates within each condition as determined from a PCA analysis of the underlying replicates. Splicing variations were assessed using the junctionCounts tool, version 0.1.2 (available at https://github.com/ajw2329/junctionCounts), against a set of splicing events that junctionCounts derived from GENCODE transcriptome version VM18. The Percent Spliced In (PSI) for each event was calculated by junctionCounts for each pseudo-replicate. The difference in PSI (ΔPsi) was calculated between each of the 9 pairwise comparisons of the 3 “Val_LNA” and 3 “Ctrl_LNA” pseudo-replicates. The results were filtered based on several parameters. 1) PSI must have been called for each pseudo-replicate using a minimum junction count (MINJC) of 15 for either the “included” or “excluded” isoform of the event. 2) The reported span of the PSI (spanPSI) call for each pseudo-replicate must not have exceeded (MAXSP) 0.25. 3) The ΔPsi between pseudo-replicates must have exceeded 0.15. To ensure consistent splicing patterns across pairs of pseudo-replicates, we calculated the “pairSum” metric, ranging from 0 to 9, which indicates the number of pairwise comparisons meeting the specified filters. Our analysis focused only on events with a pairSum of 7 or higher after filtering. To visualize the junctionCounts PSI calls for each pseudo-replicate, a UCSC genome browser track was constructed in which each splicing event is represented by two isoforms, one for the “included” and one for the “excluded” version of the event. The level of gray of the representation of each isoform is adjusted by the BED-format “score”. The score is nominally 1000*PSI for the “included” version and 1000*(1-PSI) for the “excluded” version of an event. These scores are both further reduced by a factor of (1-spanPSI) when spanPSI is non-zero, to reflect the uncertainty in the PSI call.

## ACKNOWLEDGMENTS

We thank all the members of the Sharma lab for their helpful discussions on this work. US was supported by NIH grant 1DP2AG066622-01, the Searle Scholars Program, and John Templeton Foundation award number 61748. The proteomic analysis was performed at the Vincent J. Proteomics/Mass Spectrometry Laboratory at UC Berkeley, partly supported by NIH S10 instrumentation grant S10RR025622.

## AUTHOR CONTRIBUTIONS

Conceptualization US; experimental design SL and US; methodology SL; data analysis and visualization SL, ADH, SK, and US; manuscript writing SL and US; funding acquisition and supervision US.

## SUPPLEMENTARY MATERIALs

### Supplementary Figures

**Figure S1: A)** Western blot analysis of protein isolated from caput, corpus, and cauda epididymis tissues using anti-SUMO antibody. SUMO antibody reacts with a band of the same molecular weight as the one detected using an anti-hnRNPAB antibody, indicating that the higher molecular weight band on anti-hnRNPAB Western blots represents SUMOylated hnRNPAB. The leftmost gel image is the image of the Kaleidoscope Precision Plus Protein Standard ladder from the same gel imaged under a trans-white light setting for the estimation of the molecular weight of the proteins. **B)** Western blot analysis of hnRNPAB levels in caput (CP, n=1) and cauda (CA, n=2) epididymosomes. Beta-actin was used as a loading control. **C)** Western Blot analysis using the anti-hnRNPAB antibody of proteins pull-down from epididymal cellular extract using tRFValCAC or scramble oligo. The band intensity is quantified in Figure 1F. The leftmost lane includes a protein standard used to estimate the protein sizes. **D)** Scatter plot representing all 5’tRNA fragments sequenced from small RNAs immunoprecipitated with hnRNPAB or IgG. The red circle points to tRFValCAC, which is enriched in hnRNPAB interacting RNAs. **E)** Western blot analysis using the anti-hnRNPAB antibody of proteins pull-down with tRFValCAC or scrambled oligo in DC2 cellular extract. Non-SUMOylated hnRNPAB (∼37KD) was the major form of this protein detected in DC2 cells. **F)** Quantification of band intensity of hnRNPAB in (E). **G)** Normalized read counts of tRFValCAC in DC2 cell and EVs. EVs have a significantly higher abundance of tRFValCAC relative to the cells, indicating a selective sorting of tRFValCAC in the EVs of DC2 cells.

**Figure S2: A)** tRFValCAC inhibition by Val_LNA oligo was analyzed in mESCs using the psiCHECK-2 plasmid wherein a tRFValCAC sequence was inserted into the multi-cloning site at the 3’ end of the *hRluc* gene. Val_LNA or Ctrl_LNA oligo was co-transfected with the reconstructed psiCHECK2 vector into mESCs. The transfected mESCs were then employed for the Dual-Luciferase® Reporter Assay to evaluate Renilla luciferase activity, using Firefly luciferase activity as an internal control for plasmid expression. Targeting of Val_LNA to the tRFValCAC sequence is expected to prevent Renilla expression but not Firefly expression. Renilla luciferase activity (normalized to Firefly luciferase activity) in Val_LNA transfected cells relative to Ctrl_LNA transfected cells was calculated. The plot shows the fold change of Renilla activity in Val_LNA cells relative to Ctrl_LNA cells (n=6). There was a 40% decrease in Renilla activity in Val_LNA transfected cells relative to Ctrl_LNA cells. **B)** Northern blot analysis was conducted using a probe for full-length tRNA-Val-CAC-2 in mESCs that were either transfected with Ctrl_LNA or Val_LNA. The graph shows the band intensity of tRNA-Val-CAC-2 normalized to 5S rRNA band intensity. There was no significant change in mature tRNA levels upon inhibition of tRFValCAC. **C)** Log2 fold change in the abundance of tRFValCAC in mESCs treated with RA for 120 hours relative to DMSO treatment (normalized to U6). **D)** qRT-PCR analysis showing Log2 fold change in transcript abundance of a histone gene, 2 histone modifying genes as well as a cell cycle gene in mESC treated with RA for 120 hours relative to DMSO (normalized to *B-actin*). Graphs represent data as mean ± SEM.

**Figure S3: A)** GSEA analysis showed enrichment of genes involved in mRNA splicing in the differentially expressed genes in Val_LNA embryos relative to Ctrl_LNA embryos. **B)** qRT-PCR analysis of *Srsf10* and *Srrm2* genes in mESCs treated with RA for 120 hours (to increase tRFValCAC levels). Induction of tRFValCAC resulted in an increased abundance of these two splicing regulators. **C-F)** UCSC Browser track visualization of alternative splicing events in Val_LNA and Ctrl_LNA embryos. Each splicing event is represented by two isoforms, one for the “included” and one for the “excluded” version of the event. The level of gray of the representation of each isoform is indicative of inclusion/exclusion scores, which are plotted as bar graphs shown in Figure 3. RefSeq annotated isoforms at specific exons for these genes are represented in blue.

### Supplementary Tables

**Table S1:** Table showing average emPAI value of LC-MS identified proteins in biotinylated Val-CAC oligo pulldown or scramble control oligo pulldown from epididymal cellular extract; data is ranked by t-test p-values. emPAI: Exponentially modified protein abundance index. n=4.

**Table S2:** Proteins significantly enriched in the tRFValCAC pulldown relative to scramble control and GO annotation of tRFValCAC-binding proteins in epididymis tissue. GO analysis was performed using the STRING analytical tool.

**Table S3:** Differentially expressed genes in embryos injected with Val_LNA compared to Ctrl_LNA using DESeq2 (n=15 for each condition). Cut off for calling significantly differentially expressed genes: Log2 fold change >= 1.5; *padj* value <0.05.

**Table S4:** Significant alternative splicing events (ΔPSI >0.15, pairSum ≥ 7) identified using junctionCounts. Psi is compared between each Ctrl_LNA (ctr) and Val_LNA (val) samples.

**Table S5:** Table showing % coverage and spectral counts of proteins identified by LC-MS in the pulldowns of biotinylated Val-CAC oligo or scramble control oligo from mESCs; data is analyzed using PEAKS Studio 11. n=3.

**Table S6:** Proteins significantly enriched in the tRFValCAC pulldown relative to scramble control and GO annotation of tRFValCAC-binding proteins in mESCs. GO analysis was performed using the STRING analytical tool.

**Table S7:** qRT-PCR primers used in the study. F: Forward primer. R: Reverse primer. The primers were designed using Primer-BLAST.

