## Supplementary figures and images for "A sperm-enriched 5’fragment of tRNA-Valine regulates preimplantation embryonic transcriptome and development"

### Supplementary figures

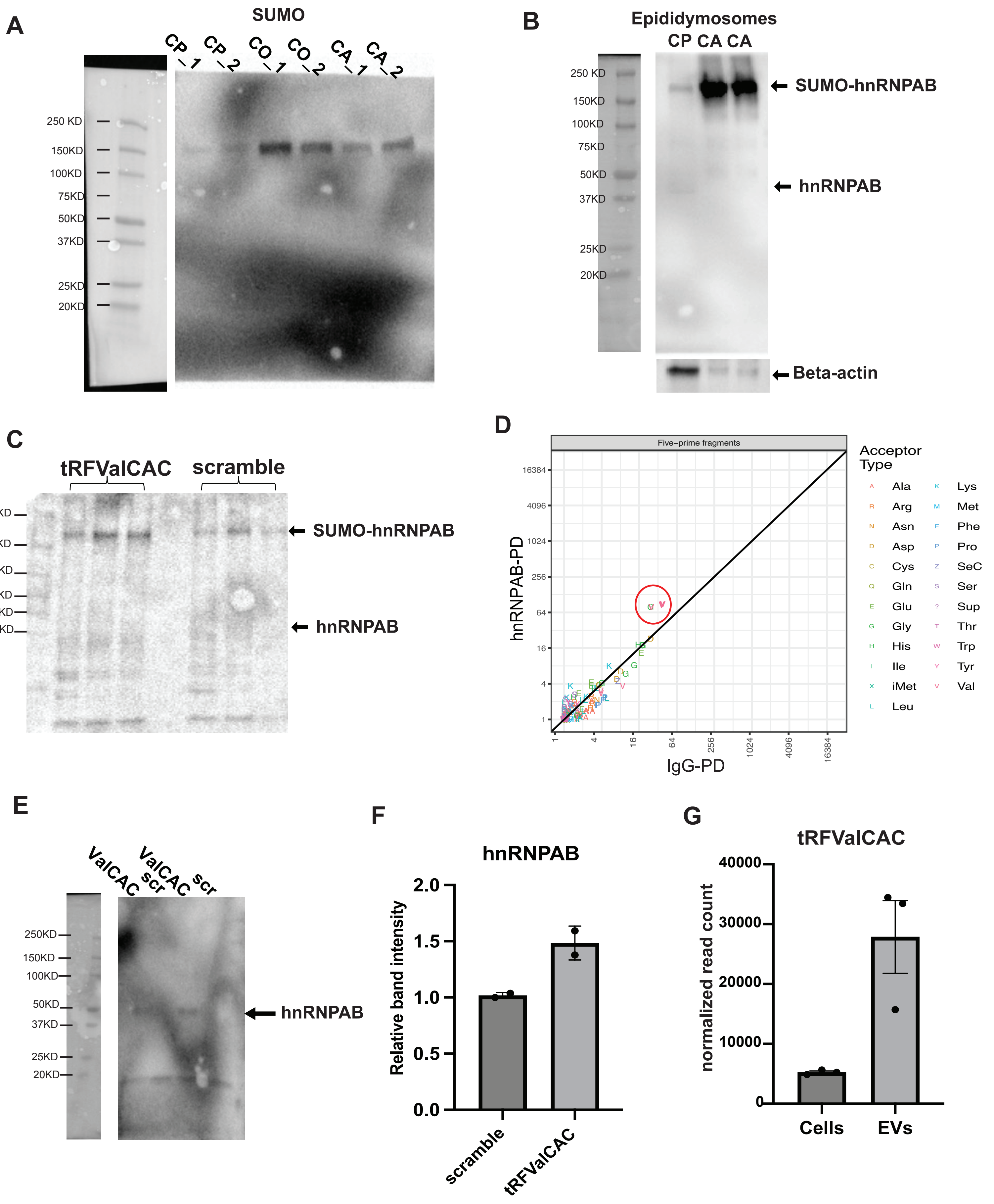

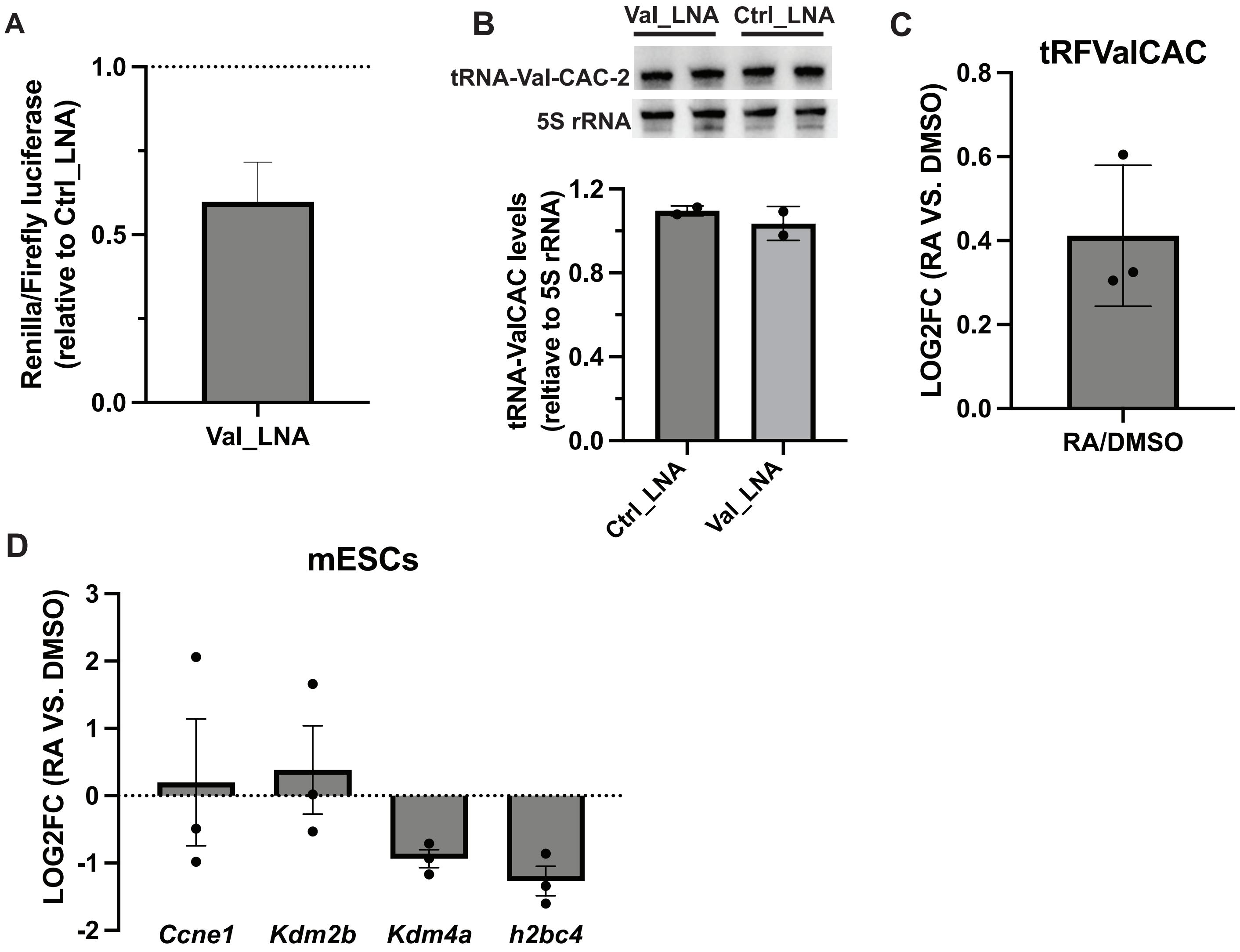

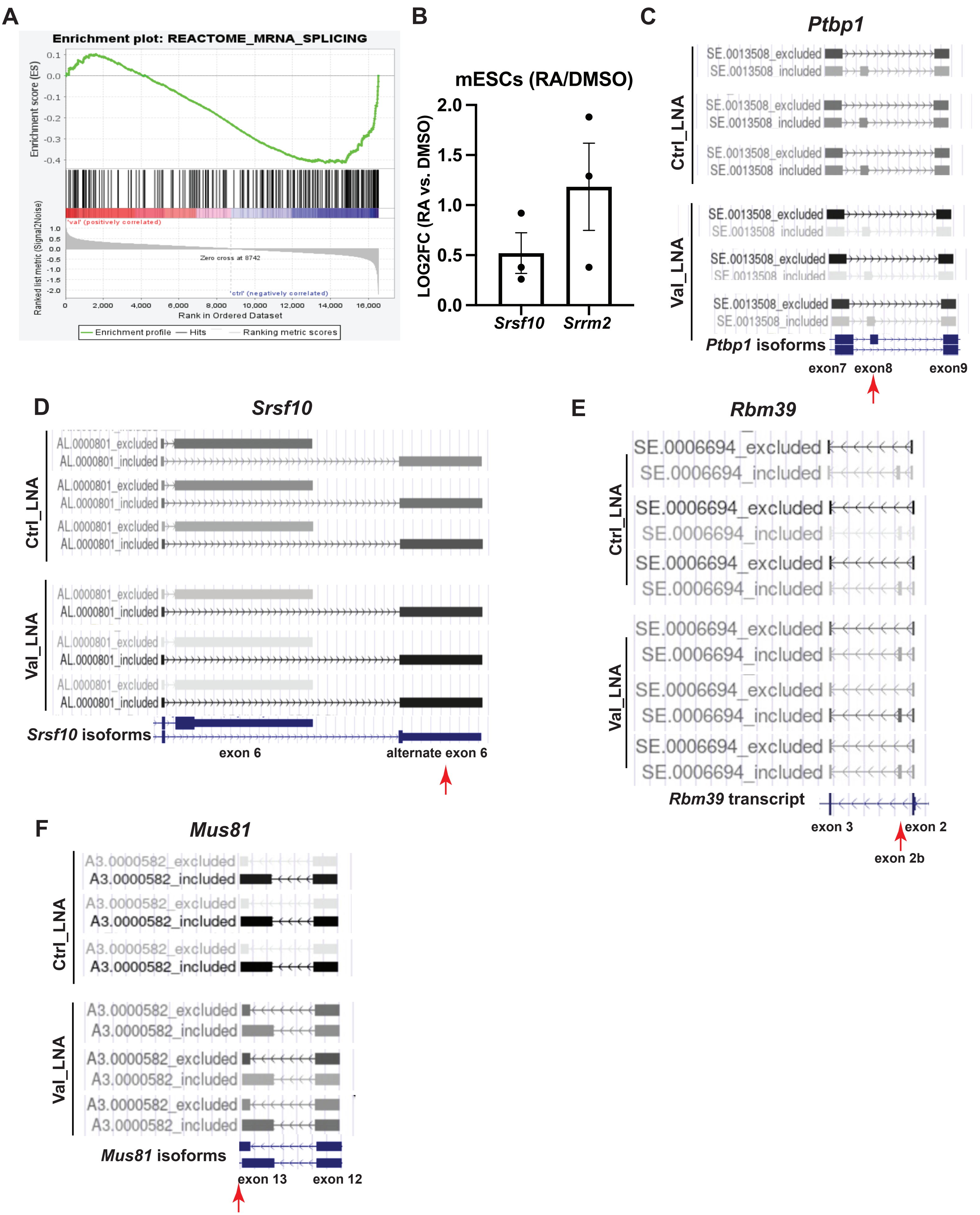
